# Distinct evolutionary trajectories in asexual populations through an interplay of their size, resource availability and mutation rates

**DOI:** 10.1101/2020.08.27.269829

**Authors:** Bhaskar Kumawat, Ramray Bhat

## Abstract

Asexually reproducing populations of single cells evolve through mutation, natural selection, and genetic drift to enhance their reproductive fitness. The environment provides the contexts that allow and regulate their fitness dynamics. In this work, we used Avida - a digital evolution framework - to uncover the effect of mutation rates, maximum size of the population, and the relative abundance of resources, on evolutionary outcomes in asexually reproducing populations of digital organisms. We observed that over extended simulations, the population evolved predominantly to one of several discrete fitness classes, each with distinct sequence motifs and/or phenotypes. For a low mutation rate, the organisms acquired either of four fitness values through an enhancement in the rate of genomic replication. Evolution at a relatively higher mutation rate presented a more complex picture. While the highest fitness values at a high mutation rate were achieved through enhanced genome replication rates, a suboptimal one was achieved through organisms sharing information relevant to metabolic tasks with each other. The information sharing capacity was vital to fitness acquisition and frequency of the genotype associated with it increased with greater resource levels and maximum population size. In addition, populations optimizing their fitness through such means exhibited a greater degree of genotypic heterogeneity and metabolic activity than those that improved replication rates. Our results reveal a minimal set of conditions for the emergence of interdependence within evolving populations with significant implications for biological systems in appropriate environmental contexts.

## Introduction

Populations of mitotically dividing cells and unicellular organisms evolve under the complex regulation of their environments. This regulation can be exerted through variations in abundance of metabolizable resources available to the population. Elegant experiments using unicellular budding yeast grown in low density sucrose-containing environments show that multicellularity can evolve under resource-poor conditions with cooperation between incompletely separated cell populations (1). Extending such observations, resource availability is theorized to have played a key role in determining the evolution of developmental mechanisms with resource-rich environments considered more ideal for the evolutionary stabilization of uniclonal, rather than polyclonal, body plans (2). Resource levels has been shown to also allow for the maintenance of cooperators within digital and experimental evolution experiments. Under resource-poor contexts, cheaters are found to dominate over cooperators (3). Whether the inverse relationship between resource level and cooperativity is altered only in the presence of cheaters is not clear.

Upper bounds on population size have also been suggested to play a contextual role in evolution: small populations are more susceptible to random genetic drift with frequent fixation of deleterious mutations. Above a certain threshold of population size, additional beneficial mutations could rescue such populations (4). On the other hand, LaBar and Adami show that that small populations show reduced susceptibility to drift, and hence their genotypes are less likely to acquire small-effect deleterious mutations (5). Another exploration of this problem under a game theoretic framework suggests that small population sizes consisting of participants that can utilize memory on longer time scales can evolve to cooperate (6).

The role of mutation rates in evolution of asexual populations has been theoretically explored by several workers (7–9). Individual mutations may have a deleterious, neutral or beneficial effect on the fitness of the populations within which they arise; fitness-enhancing mutations constitute only a small proportion within this possible set (10). However, the rate of adaptation has been shown to increase under high mutation rates (11), but this is not without a limit: very high mutation rates can lead to a catastrophic decline in fitness (mutational meltdown) (12). Increased mutation rates within heterogeneous populations result in evolution of populations to lower fitness while attaining higher robustness: a hypothesis known as the “survival of the flattest” (13,14). Further, in experimental models, increased mutagenesis has been shown to be a genotypic response to environmental stresses (15–17) and can in turn increase evolvability (18).

Organisms inhabiting the natural world evolve within their diverse and often variant environments, with all the above factors simultaneously constraining and biasing their phylogeny. In this article, we examine the interplay between the three factors - mutation rate, resources availability and maximum population size and ask whether these factors amplify or offset each other’s effects in determining the likelihood of populations evolving through specific strategies. To achieve this, we utilize the Avida artificial life platform, wherein mutating, reproducing, and resource-metabolizing digital organisms are allowed to evolve under distinct values of these environmental variables. We analyze the constitution and frequency of predominant genotypes emerging from the evolutionary runs of ancestral organisms that begin with a basic reproduction program and the ability to perform a simple metabolic task. Our studies shed light on the necessary and sufficient conditions for populations to evolve and improve fitness through inter-independent and dependent mechanisms (with distinct genotypic underpinnings), under mutagenesis and environment-permissive contexts.

## Methods

### Avida computational model

The Avida digital evolution platform is a computational framework that allows simulation of self-replicating organismal populations in a lattice-based virtual world (19). It has been used previously to study evolutionary dynamics of multiple systems in order to answer fundamental questions in evolution and ecology (14,20–22).

Each organism in Avida consists of the following components,

a. Three registers that can store a 5-bit number (the unit of information in Avida) each and are individually accessible by the organism
b. Two stacks that can store any number of 5-bit numbers in a first-in-last-out fashion, and
c. A CPU that executes the instructions that form the *genotype* of the organism (the software that runs on the above hardware).

A Turing-complete set of genomic instructions allow these organisms to copy the genome, replicate by division, perform logical/mathematical operations, control the flow of execution (using jumps and loops), perform input/output to interact with the environment or send information to neighbors. The environment provides 5-bit numbers as metabolic substrates that the organisms can read and manipulate in order to generate a resultant output. If the output maps to the inputs by one of the tasks rewarded in the environment, the organisms take up a certain fraction of an external *resource* (implemented as a global chemostat that maintains a certain level of resources in each lattice-site in the world) and are rewarded a corresponding *merit*, a quantity that essentially determines the speed with which an organism genome is executed. In turn, genomes that execute faster, copy faster and give rise to more offsprings per unit time. To accomplish reproduction, however, they also need to have a working copying-and-division mechanism that is implemented in the ancestral seed genome as a *copy-loop*. This ancestral copy-loop has two main functions - (a) To copy the parent genome instruction-by-instruction until the entire genome has been copied, and (b) to divide the offspring only after the copying is complete, and place this offspring in a new neighboring world cell.

Apart from sending generated task results to the environment for verification (using output instructions), the organisms are also able to send the values stored in their memory to their eight nearest neighbors by executing a set of instructions. These *messaging* instructions can either send intermediary metabolic information (for example the result of an unfinished numerical algorithm) or allow inter-cellular regulation by acting as a signal for other genomic instructions. The organisms have a faced direction in the world and can *rotate* in order to choose either the receiving neighbor (for messages) or to choose the direction in which to place an offspring.

The organisms are thus free to innovate in three major areas - metabolism, reproduction, and cooperation. The set of instructions that are relevant in our study and allow the organisms to innovate these mechanisms is given in Table 1. There are a total of nine tasks available in the environment, each with a corresponding resource. The list of these tasks and their merit payoffs are given in Table 2.

**Table 1.**
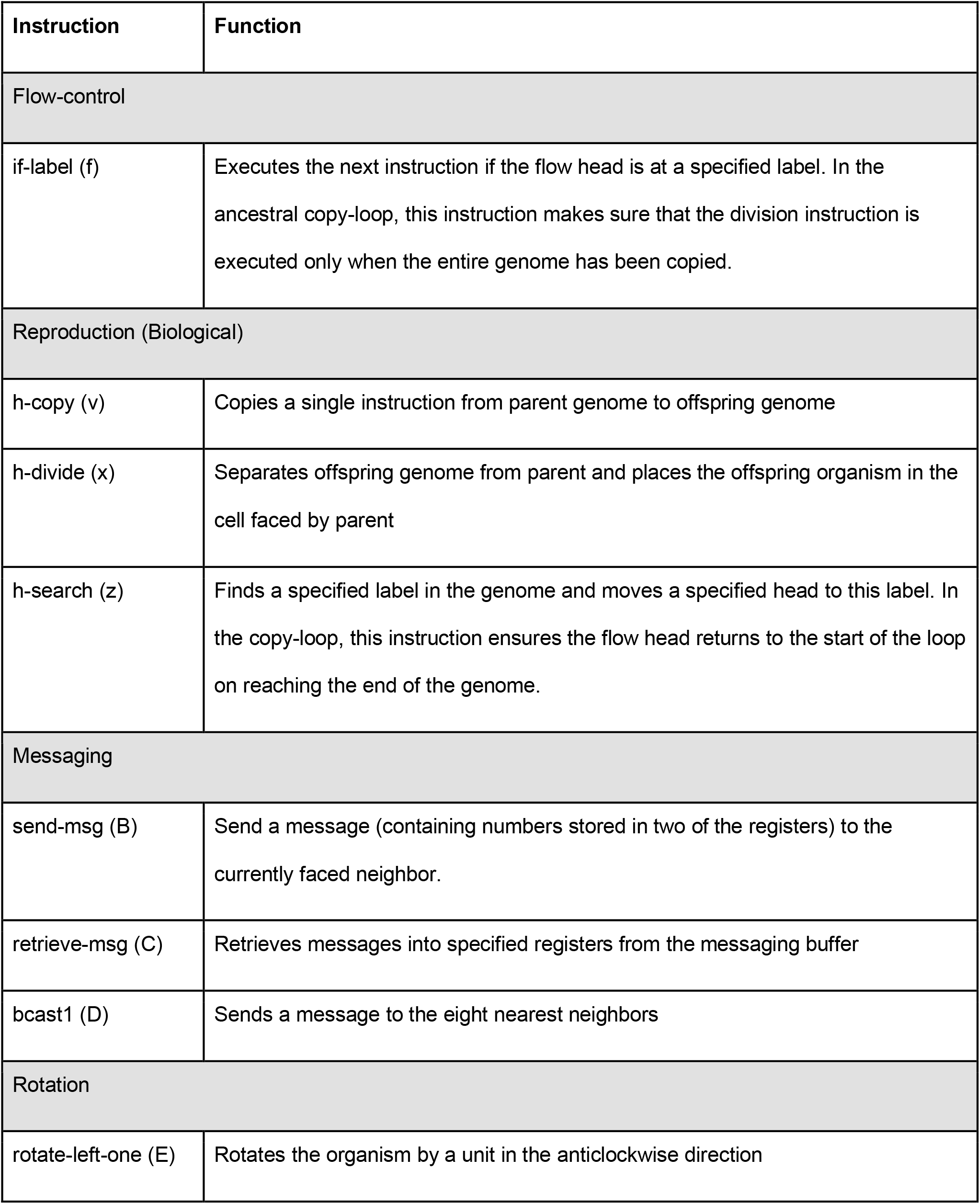
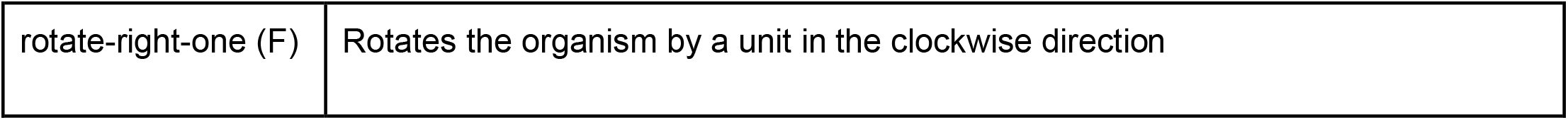
Major instructions of interest in Avida and their functions.

**Table 2.** Metabolic tasks in the environment and their merit payoffs. Adapted from (3)

| Task | Operation | Merit<br>reward |
| --- | --- | --- |
| NOT | $\sim A$ | 1 |
| NAND | $\sim (A \wedge B)$ | 1 |
| AND | $A \wedge B$ | 2 |
| ORN | $A \vee \sim B, \sim A \vee B$ | 2 |
| OR | $A \vee B$ | 4 |
| ANDN | $A \wedge \sim B, \sim A \wedge B$ | 4 |
| NOR | $\sim (A \vee B)$ | 8 |
| XOR | $(A \wedge \sim B) \vee (\sim A \wedge B)$ | 8 |
| EQU | $(A \wedge B) \vee (\sim A \wedge \sim B)$ | 16 |

### Experimental Setup

For all the experiments presented here, a single ancestral organism is released in an Avida world (with periodic boundaries) and allowed to reproduce to give rise to a population that evolves over 100,000 updates, the basic unit of time in Avida. Simulations are run with two values of the mutation rate, at different levels of resource availability, and population size conditions. The ancestral genotype consists of a simple copy-loop with a primitive task definition that allows it to perform the simplest task in the environment. It also has a single rotation instruction, which allows the organism to change the direction it faces in the world and thus can facilitate messaging and/or dispersal of offsprings after division.

Under the normal (or high as mentioned in the paper, relative to low below) mutation rate, there is a 0.75% probability of a genomic instruction being substituted with another on divide. Under the low mutation rate, substitutions are made with a probability of 0.075%, an order of magnitude lower than normal. The sequence length is restricted to 120 and mutations events are limited to substitutions to conserve this length.

The resource availability is varied between three values, each an order of magnitude higher than the previous one. The lowest of these is at an absolute value of 10,000 and is chosen such that there are no secondary effects that restrict proper mutation and selection over the duration of the experiments (such as a loss of viability due to starvation). The upper bounds on population size are varied between 100, 200 and 500. Each treatment is repeated for 50 replicates.

### Fitness distribution analysis for major innovations

Avida has an internal measure of fitness that is calculated by taking a ratio of the metabolic rate (i.e. merit) to the gestation time of an organism (time required for replication from the start of genome execution). However, both these values are calculated with the organism in isolation and thus the definition does not account for cooperative benefits to fitness. To remedy this, we utilize a slightly modified metric that places 200 copies of the organism in a world, lets the population stabilize for a hundred updates and then calculates the average number of births per update for the next 400 updates. The mutation rate is set to zero during the entire process and thus prevents the sequences from evolving.

Each mutation rate has its own set of major innovations that appear on the fitness distribution (as discussed in Results). To generate this fitness distribution, we collect the genotypes from all the treatments and replicates under a mutation rate and plot the kernel density estimate for the fitness values. The high-density peaks that appear are taken as representatives of different mechanisms that appear with a high probability. As discussed later, resource levels and maximum population size do not alter the position of these peaks but rather determine the probability of finding a genotype within these states (Figure 1).

**Figure 1.**
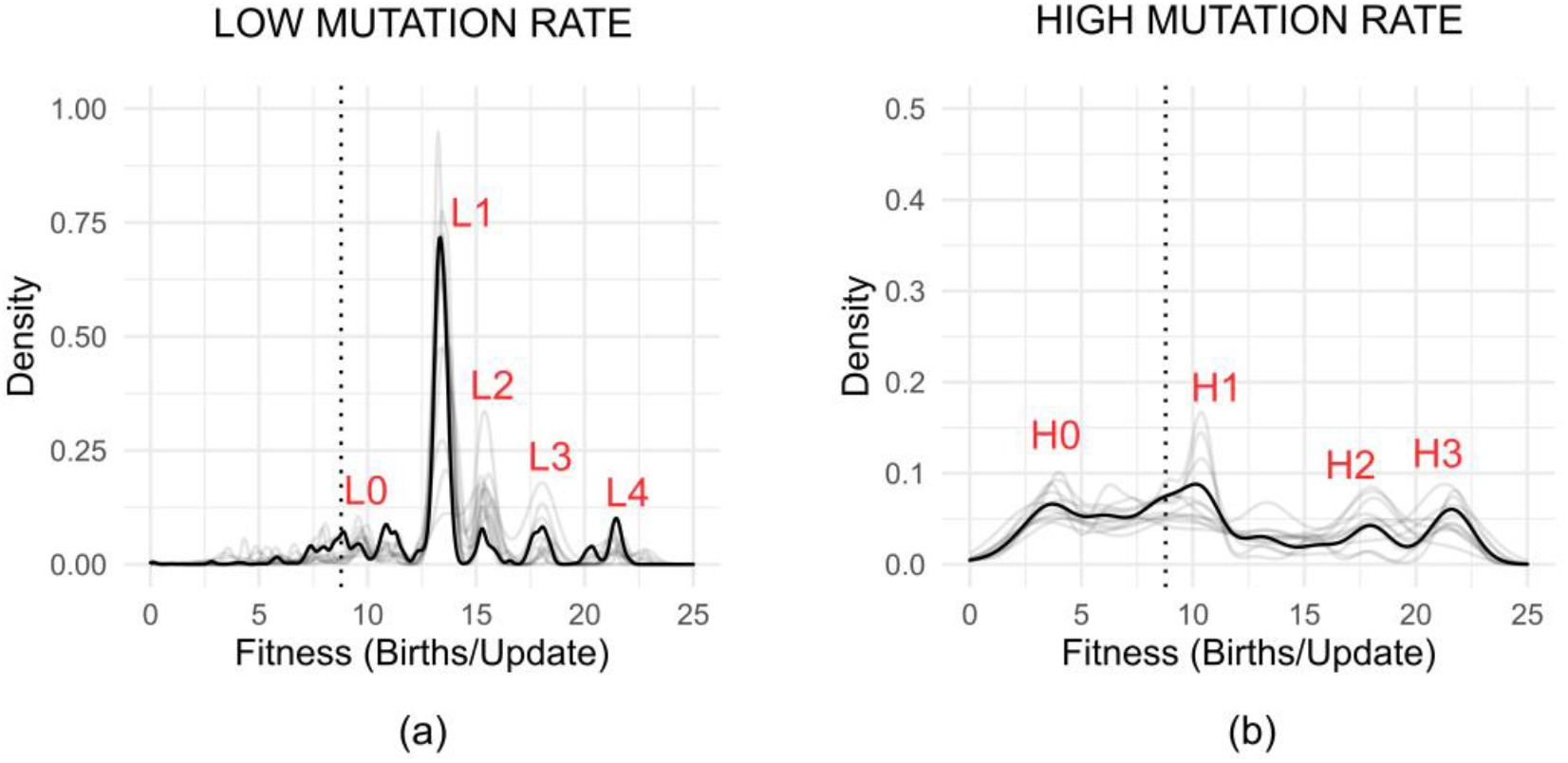
Fitness distribution of genotypes obtained from simulation at low and high mutation rates of a population of an ancestral organism (Dark line: resource levels of 10k and population size of 500; Light lines: all other combinations of these variables, see Supplementary Figure 2). (a) At low mutation rate, five major peaks (labeled L0-4) are obtained in the distribution. (b) At a high mutation rate, the distribution becomes more heterogeneous and four major peaks (H0 to H3) are observed. The dotted vertical line for both spectra represents the fitness of the ancestral organism. Y axis represents density (calculated from a kernel density estimate with area under the curve normalized to 1) and X axis represents fitness (defined as the number of births/updates)

The genotypes from each peak are sampled by considering a narrow region around the peak maxima (within 2xKDE bandwidth). These sampled genotypes are then analyzed for common features to deconvolute the underlying mechanisms that give rise to the corresponding peaks.

### Marginal Utility of a Process

We define a metric called the marginal utility of process for each genotype given a process that it can possibly use to acquire fitness. This metric is calculated as follows.

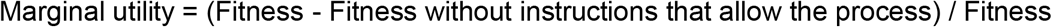

Fitness without the process instructions is calculated by substituting all the relevant instructions in the genotype sequence with a null instruction and measuring the fitness of the resultant sequence.

### Heterogeneity Calculation

Evolved populations (i.e. each replicate) are classified into one of the major peaks found at that mutation rate by noting down the position of the population’s dominant genotype on the fitness distribution. Following this, we calculate the per-site entropy for the set of genomes in each of these populations and sum these values for all 120 sequence sites to get the population heterogeneity. For each position *i* in the genotype sequence, the per-site entropy is given by,

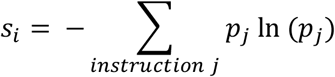

Where *p*_*j*_ is the probability of instruction *j* appearing at position *i* in the sampled genotypes.

### Software

Version 2.14.0 of Avida was used for these simulations. Automation of the analysis, data retrieval and cleanup were done using Python version 3.8.2 (with numpy 1.19.0 and matplotlib 3.2.2). Data analysis and plotting was performed using R version 4.0.2 and the ggplot2 library (version 3.3.2). All these experiments were conducted on x86-64 machines running GNU/Linux kernel 4.15.0.

## Results

### Mutation rate regulates diversity in fitness acquisition and associated genotypes

We began by comparing the genotypes obtained after 100,000 updates of evolving the ancestral Avida organisms and plotted the fitness spectra for genotypes obtained at high and low mutation rates (and at three resource availability values and three population sizes) in Figure 1. For the low mutation rate, five major peaks were observed in the fitness distribution (Figure 1a, peaks denoted as “L0”, “L1”, “L2”, “L3” and “L4”). The peaks were found to be progressively higher in fitness than that of the ancestral organism (denoted by a vertical dotted line). For the high mutation rates, we observed four major peaks (Figure 1b, denoted “H0”, “H1”, “H2”, and “H3”). Peak H0 was lower in fitness and peak H1, H2 and H3 were progressively higher than the ancestral type. We note that only a single dominant peak was observed for most runs (Supplementary Figure 1). Therefore, each of the fitness peaks likely represented a stable state and a population evolved stochastically into one of these states. In summary, the mutation rate was the primary determinant of the set of major genotypic fitness values that emerge at the end of evolutionary runs. The resource availability and population size only affected the relative probability of genotypes attaining one of these values (Supplementary Figure 2). The representative genotypes were isolated from a small region around the peak maxima as given in Supplementary Table 1.

### Genomes that evolved at a low mutation rate diversify by differences in genome replication rate and metabolism

We examined the genotypes cognate to the fitness peaks to identify the differences in sequence that accompanied their evolution. The predominant difference between the genomes from the corresponding peaks (evolved at low mutation rate) was found in their copy-loops (Figure 2a): the h-copy (v) instructions were numerically distinct in the copy-loops of genomes across the peaks.

**Figure 2.**
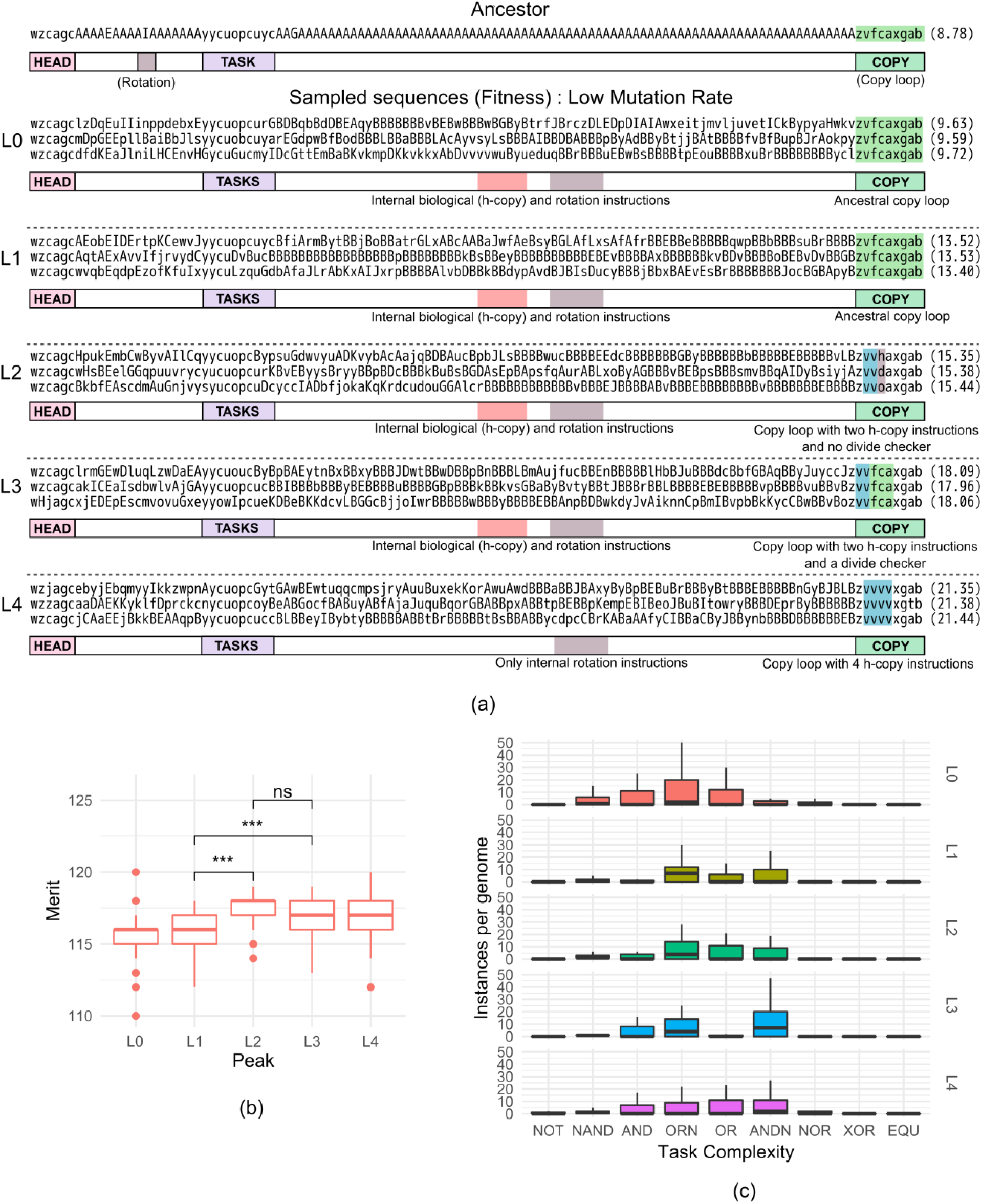
(a) Genome sequences and schematic depiction of key genomic elements from the ancestral organism (top) and three representative organisms from L0-L4 peaks obtained from simulations at the low mutation rate (see Figure 1A). The copy-loop (highlighted in green) is the major source of difference between these sequences. Fitness for the sequences are provided (b) Box plots depicting merits (speed of the organism’s virtual CPU, a proxy for metabolic activity of the genotype) of 100 genotypes sampled from each of L0-L4 on the fitness distribution of simulations performed at a low mutation rate. Genomes from L0 and L1 are metabolically less active compared to L2, L3, and L4 (statistical significance measured using the non-parametric Wilcoxon test; ***: p<0.001). (c) Task complexities for genomes (estimated roughly by the number of “NAND” instructions required to perform a task, increases from left to right) from L0-L4 peaks on the fitness distribution obtained from simulations at a low mutation rate. Higher fitness peaks are found to perform more complex metabolic tasks. The Y axis represents the number of times that a genotype belonging to these peaks performs a task over a single execution of the genome.

Organisms from peaks L0 and L1 showed a copy-loop identical to the ancestor, containing a single h-copy (v) instruction and an if-label instruction (f) that validated the completion of the genome duplication process and only then allowed the offspring genome to segregate from the parent. Organisms in L2 used two h-copy instructions which allowed them to copy two genomic instructions every iteration, but did not have a validation system, potentially allowing the birth of partially copied non-viable genotypes. The overall effect was an increase in fitness, but organisms with these genotypes nevertheless had a lower fitness relative to the ones that retain the validating if-label instruction as found in L3. L4 genomes (which showed the highest fitness) were able to evade this requirement by incorporating a large number of h-copy instructions (generally four), essentially allowing them to rapidly copy the entire genome without offspring sequence validation. The low mutation case thus contrasts the differences in genome-copying robustness that evolves under different conditions.

In addition to copy-loop differences, we observed the duplication of elements present within the copy-loop and their transition to the non-copy-loop regions of the genotypes for all peaks except genomes of peak L4. However, measuring the marginal utility of these *internalized* elements showed that they did not contribute significantly to the fitness of the genotypes (Supplementary Figure 3; median values for all L-peaks were found to be lesser than 0.05, indicating that the addition of these internalizations added less than 5% fitness advantage to the sequences).

The organisms from different peaks were also observed to exhibit differences in their capacity for utilization of resources (metabolism). L0 and L1 organisms acquired a lower cumulative median merit than those from peaks L2, L3, and L4 (Figure 2b). The complexity of tasks performed showed a progressive increase across organisms from peaks L0 to L4 (Figure 2c).

These results led us to investigate whether changes in the copy-loop contributed to determination of fitness and merit for the evolved genotypes. Replacement of the copy-loop in a representative genome from peak L0 with a copy-loop containing multiple h-copy instructions (from a L4 genotype) instantaneously increased the fitness to an intermediate value (Supplementary Figure 4). Allowing its population to evolve accentuated the fitness to L4 fitness levels. The merit on the other hand, increased instantaneously to the L4 value, confirming that the nature of the copy-loop determines the metabolic capacity of the organisms. A reverse experiment (transplant of L0 copy-loop into L4 genomes) also showed a similar reliance of fitness and merit on the reproductive region of the genome (Supplementary Figure 5). Specific genomic ‘chimerization’ influenced the viability of the organisms: although all L4 organisms with L0 copy-loops were viable, addition of L4 copy-loop to L0 organisms instantaneously brought down the viability by 80%.

### Genomes with high mutation rate diversify through differences in evolutionary mechanisms

In keeping with the analysis above, genotypes associated with the four fitness peaks observed for evolution under a high mutation rate were analyzed for inter-sequence differences (Figure 3a). Peak H0 genotypes showed a dismantling of the ancestral copy-loop and the internalization of copy-loop elements (as was also observed in the fitness peaks of the low mutation rate runs). The change in fitness that two sample genotypes from this peak experienced over evolutionary time showed that the decline was concomitant with the occurrence of a mutation that removed the h-search instruction (Supplementary Figure 6a). In spite of the breakdown of this vital copy-loop, these organisms survived by accommodating internalized copy-loop elements in the interior of their genome (removal of such copy-loop elements led to loss of fitness for these genotypes; Supplementary Figure 6b). Interestingly, copy-loop element internalizations precede the dismantling of the ancestral copy-loop, suggesting that the former is an inherent feature of genomic evolution (it is seen for all fitness peaks except for the highest one under low mutation rate) that is ‘exapted’ to prevent extinction in the contingent case of copy-loop loss (23).

**Figure 3.**
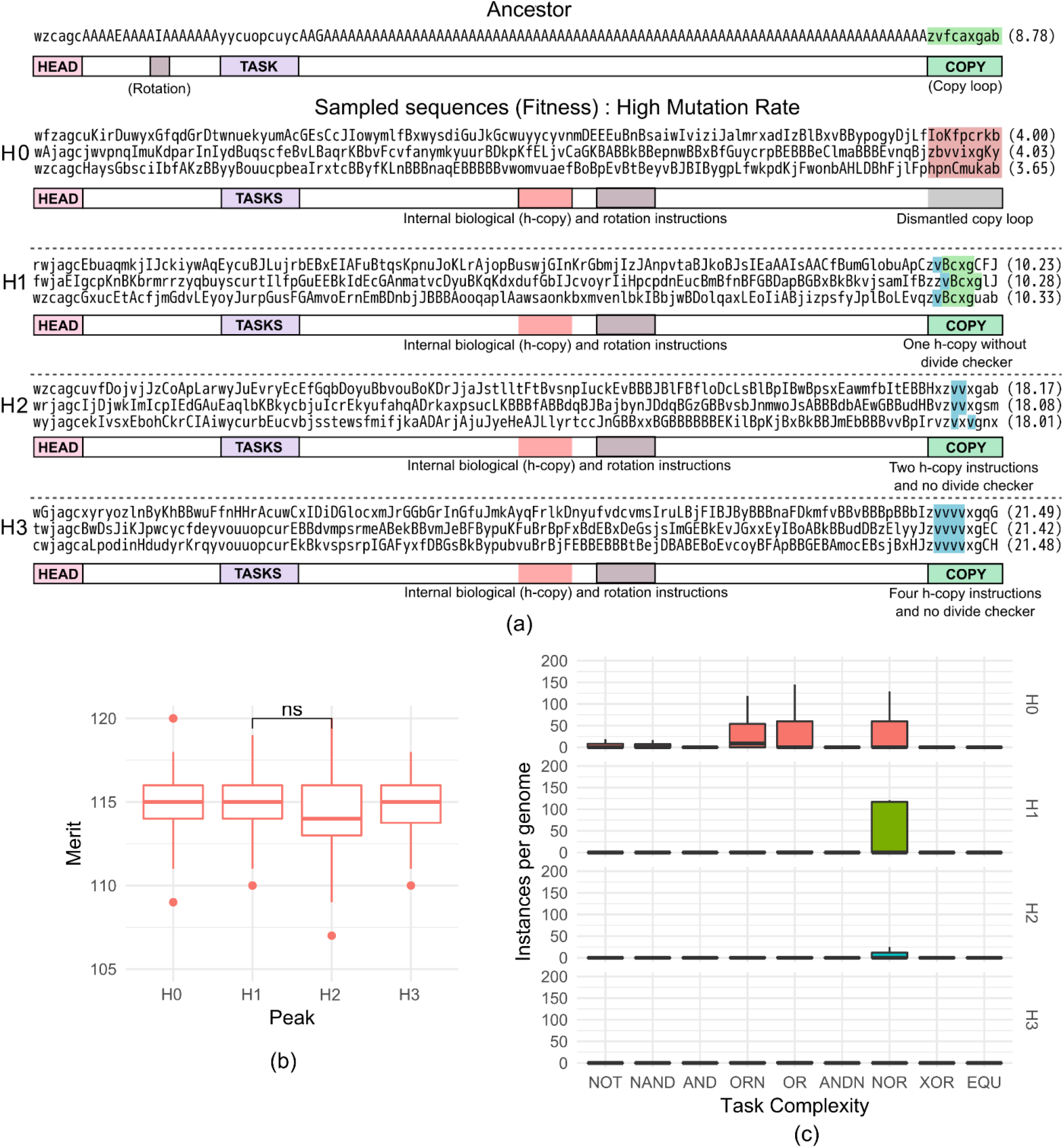
(a) Genome sequences and schematic depiction of key genomic elements from the ancestral organism (top) and three representative organisms from H0-H3 peaks obtained from simulations at the high mutation rate (see Figure 1B). The copy-loop is the first point of difference between these sequences. H0 genotypes have a copy-loop devoid of the ancestral copying functionality. Fitness for the sequences are provided. (b) Box plots depicting merits (speed of the organism’s virtual CPU, a proxy for metabolic activity of the genotype) of 100 genotypes sampled from each of H0-H3 on the fitness distribution of simulations performed at a high mutation rate (statistical significance measured using the non-parametric Wilcoxon test). (c) Task complexities for genomes (estimated roughly by the number of “NAND” instructions required to perform a task, increases from left to right) from H0-H3 on the fitness distribution obtained from simulations at high mutation rate. The Y axis represents the number of times that a genotype belonging to these peaks performs a task over a single execution of the genome.

H1 genotypes retained a copy-loop consisting of only the basic replication instructions (h-copy and h-divide) required for copying but did not implement an if-label instruction that allows detection of complete copying. Genotypes from H2 and H3 showed progressively greater h-copy instruction numbers in their copy-loop suggesting their higher fitness was a result of an enhanced rate of replication. Metabolically, peak H0, H1, H2 and H3 genomes were found to be almost equivalent in merit to each other ^1^ (Figure 3b). H0 and H1 performed a very large number of metabolic tasks per cycle (compared to the genomes obtained at a low mutation rate). In H2 and H3, the large number of executions were replaced with high reward (and high complexity) tasks (Figure 3c).

We observed that in addition to changes in copy-loop sequences and metabolic task elements, messaging (information sharing) instructions were also characteristic of genotypes representative of peak H1, H2 and H3. On sampling and testing the utility of messaging in these genotypes (see Methods), we found that messaging contributed significantly to the fitness acquired by genotypes associated with peak H1, but not H2 and H3 (Figure 4). Interestingly, the increase in fitness over the ancestral type for these (H1) genomes was approximately equal to the contribution from messaging (∼20%).

**Figure 4.**
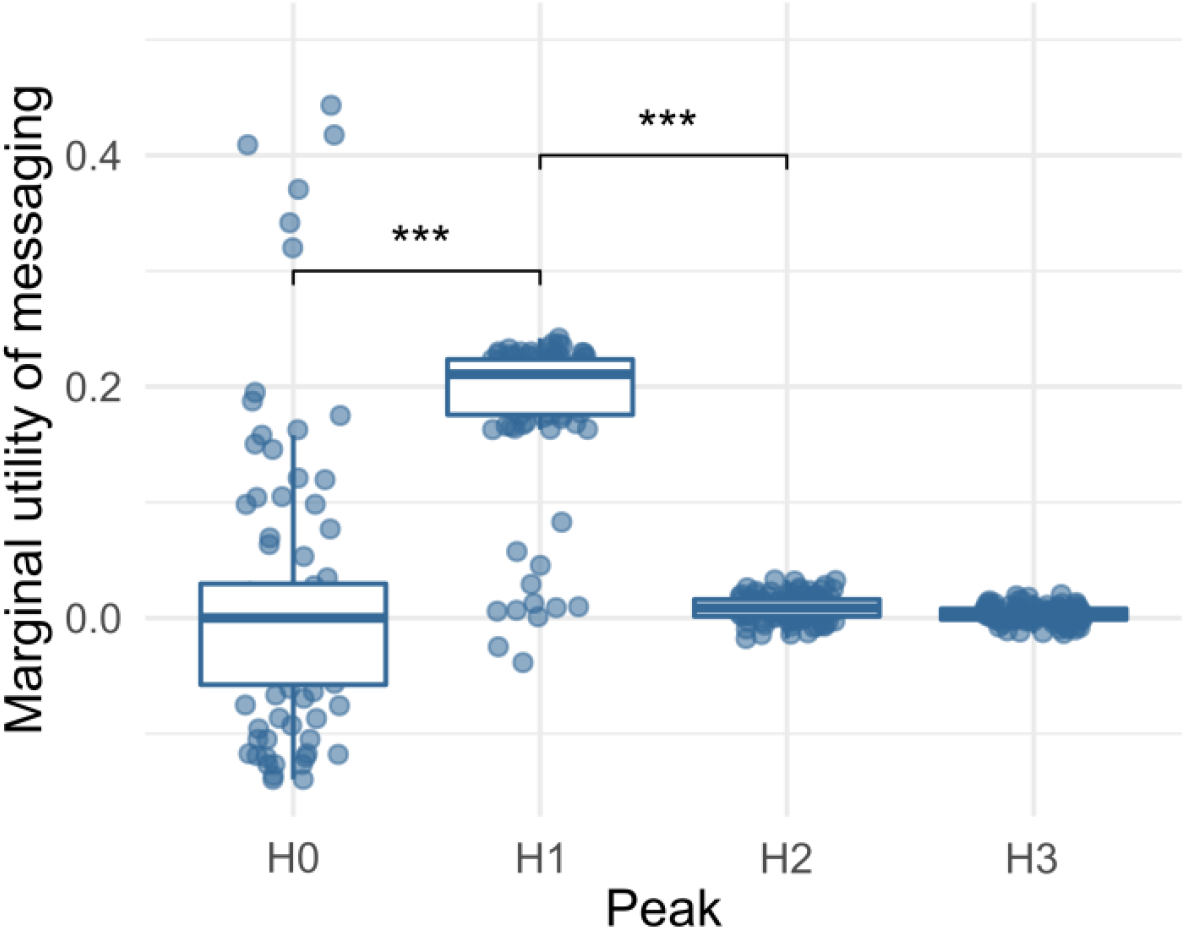
Box plot showing marginal utility of messaging for genomes (fractional addition to fitness provided by messaging, see Methods) sampled from peaks H0-H3 of the fitness distribution obtained from simulations at a high mutation rate (see Figure 1B). Only peak H1 utilizes messaging as a facilitator for acquiring a higher fitness (significance measured using the non-parametric Wilcoxon test ***: p<0.001).

When the copy-loop of H0 genotypes was replaced with that of a peak H3 genotype, and the hybrid genotype was allowed to evolve, we did not observe an instantaneous increase in fitness: rather, the initial unchanged median fitness gave rise to rise to multiple genotypes both lower and higher in fitness than the initial hybrid (Supplementary Figure 7). H1 genotypes with their copy-loops replaced by H3 copy-loops showed an instant increase in fitness which diversified over evolutionary time (Figure 5a). The viability, just as in the low mutation rate case, was affected as a result of the change and only 20% organisms survived to evolve. Removal of the cognate copy-loop from H1 and replacement with H3 copy-loop dropped the marginal utility of messaging to 0 (Figure 5b). Consistent again with the low mutation rates, 100% of peak H3 genotypes, upon their copy-loops being replaced by H1 counterpart, evolved to fitness levels congruent with H1 genotypes (Figure 5c). To our surprise, an increased marginal utility of messaging was found to be conferred by the addition of the H1 copy-loop to genotypes from H3 (Figure 5d). Our results indicated that the copy-loop played a major role in determining the benefits that an organism accrues through messaging.

**Figure 5.**
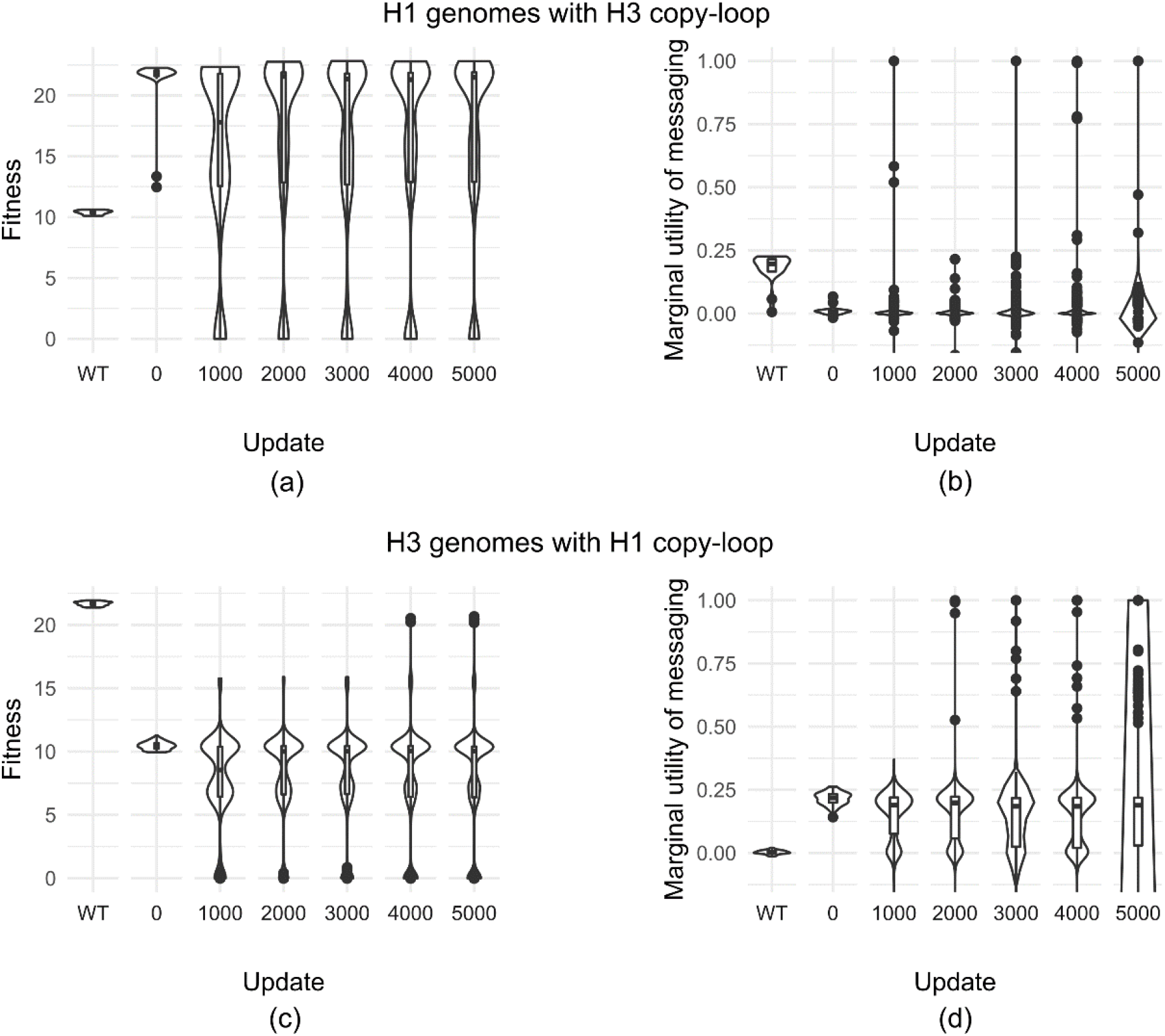
Violin plots showing (a) fitness and (b) marginal utility of messaging of genomes sampled from H1 upon their endogenous copy loop being replaced with H3 copy-loop and evolution for 5000 updates. (c) Fitness and (d) marginal utility of messaging of genomes sampled from H3 upon their endogenous copy loop being replaced with H1 copy-loop and evolution for 5000 updates. Updates on the x-axis represent the evolutionary time for which the hybrid genotypes were evolved. “WT” denotes these measures for the recipient genomes without copy-loop replacement.

#### Populations dominated by cooperative genotypes are more genetically heterogeneous

We asked whether the fitness values associated with peak H1 genotypes were suboptimal with respect to H2 and H3 counterparts, so as to accommodate greater genetic heterogeneity within the populations when they arose. Shannon entropic measurements made across sequence alignments of populations that evolved under high mutation rates showed that H1 populations were genetically more heterogeneous compared with populations from the other peaks associated with high mutation rate (Figure 6). This variation was seen to be almost invariant across resource levels and population sizes (except at the lowest population size and 100k resource levels where the heterogeneity was found to be lower; Supplementary Figure 8). To check whether this difference in genotypic heterogeneity was specific to the mechanism of fitness acquisition, we transplanted copy-loops from H1 and H3 into each other’s genotypes. To ensure that the process of transplant itself doesn’t introduce changes in heterogeneity, we performed a control run transplanting H1 populations with a H1 copy-loop (Supplementary Figure 9). Addition of H3 copy-loop to H1 populations decreased the population heterogeneity to H3 levels over 5000 updates of evolution (Figure 6b). A reverse experiment of replacing H3 populations with H1 copy-loop significantly increased the population heterogeneities as well (Figure 6c).

**Figure 6.**
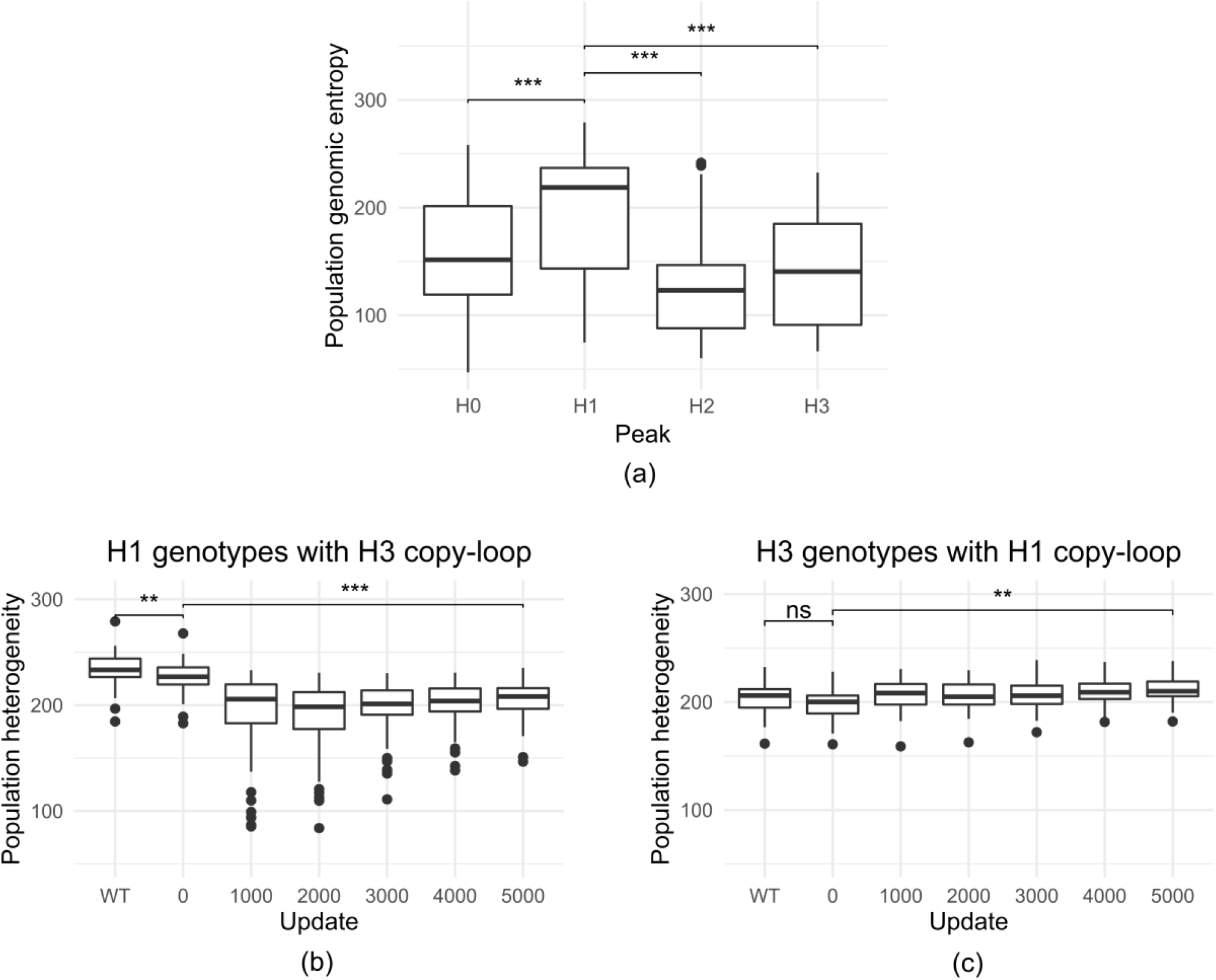
(a) Box plots showing genomic heterogeneity within populations belonging to H0-H3 peaks of the fitness distribution obtained from simulations at a high mutation rate (see Figure 1B). (b) Change in population heterogeneity when populations dominated by H1 are transplanted with H3 copy-loop and evolved for 5000 updates (Populations evolved under maximum size 500 and 1000k resource availability). (c) Change in population heterogeneity when populations dominated by H3 are transplanted with H1 copy-loop and evolved for 5000 updates. Population heterogeneity is measured by calculating the sum of per-site genomic entropies (See methods, significance calculated using the non-parametric Wilcoxon test; **: p<0.01, ***: p<0.001). “WT” denotes heterogeneity of the recipient population without copy-loop replacement.

### Maximum population size and resource levels regulate frequency of the occurrence of distinct fitness peaks

In order to quantify the extent to which resource availability and maximum population size affect the probability of appearance of a particular mechanism, we isolated genotypes from within a narrow range around the major peaks (see Methods) and calculated their frequency within the total number of genotypes. This allowed us to quantify the major trends that we observed qualitatively in Supplementary Figure 1. We saw a progressive decrease in the frequency of occurrence of L1 genotypes both when resource levels and population size were increased (Figure 7a) under low mutation rates. The frequency of occurrence of peak L0 and L4 fitness remain unchanged. Peak L2 showed mixed trends whereas L3 showed an increase with increased resource levels and population size.

**Figure 7.**
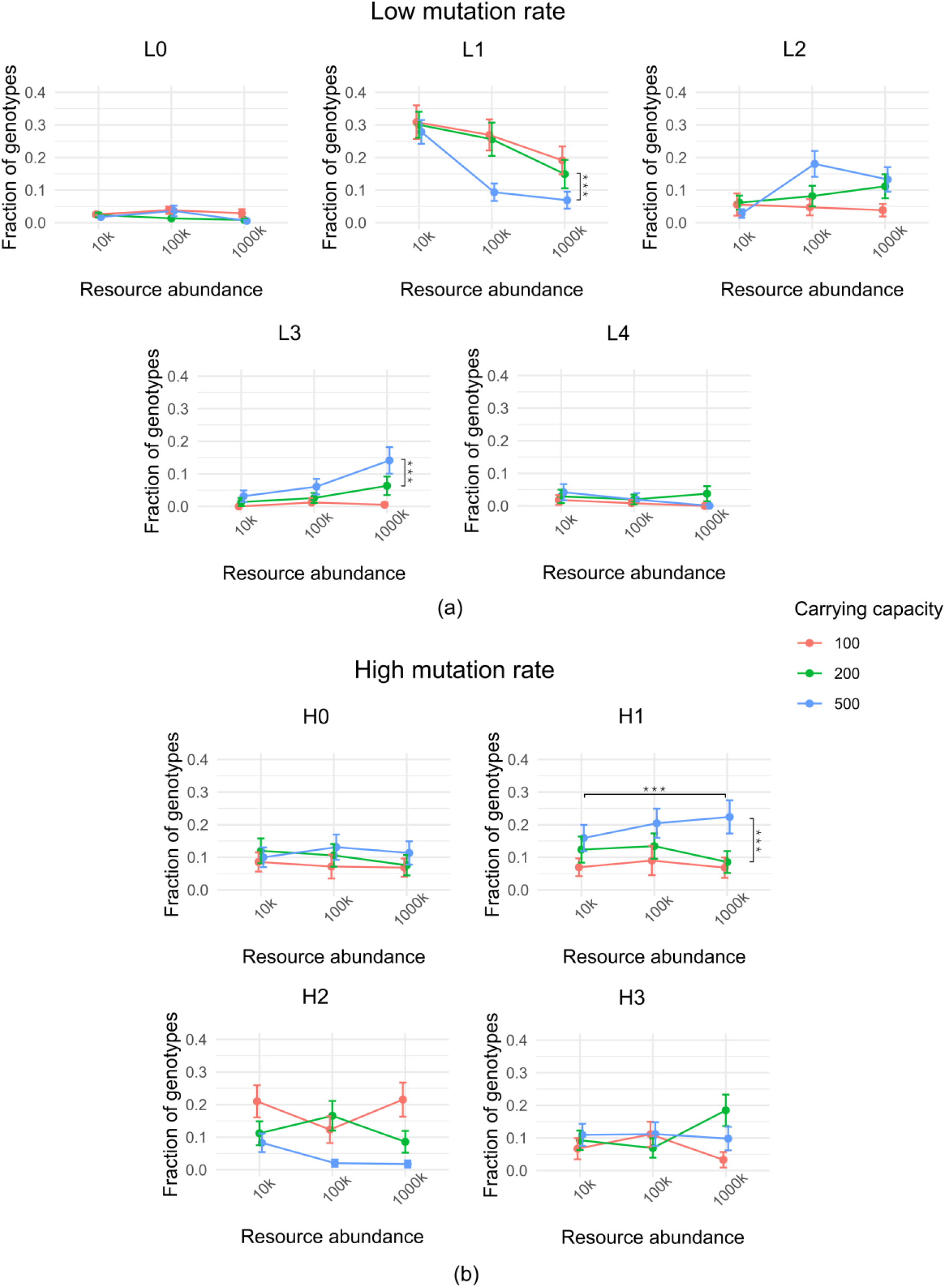
Graphs showing fractional proportion of genotypes of peak L0-L4 (a) and H0-H3 (b) obtained from the fitness spectra from simulations run at low- and high-mutation rates respectively (see Figure 1A and 1B) at 3 resource levels (10k, 100k, and 1000k) with population size limits of 100 (orange), 200 (green) and 500 (orange). (Bootstrap analysis with a sampling size of N=30 out of 50 populations resampled 1000 times, significance measured using an unpaired t-test ***: p<0.001)

The frequency variations presented a more complex picture for evolution under high mutation rates. Whereas H0 frequencies were unaffected by the two environmental factors, those for H2 and H3 did not show clear patterns (Figure 7b). On the other hand, patterns were discernible for the peak H1 frequencies: its occurrence increased with higher population sizes for all resource levels. Additionally, increasing resource levels also accentuated H1 frequency but only at the highest population size (500). The highest frequency for H1 was seen for evolutionary runs where the maximum population size and the resource levels were highest.

## Discussion

In this work, we have explored the interplay between the environment and the driving force of phenotypic variation: mutation rate, in the exploration of an adaptive landscape by a population of digital organisms. It is increasingly being recognized that the effect of mutation rate on fitness needs to be examined in the context of the environment in which the evolution takes place. In this manuscript, we consider two environmental parameters: maximum population size and resource levels. A positive regulation effect of population size on fitness can be explained by the higher probability of mutants appearing in the population; in fact under high mutation rate, bigger population size allow for greater exploration and fixation of genotypes that may otherwise never have been explored (24). Resource availability, on the other hand has been shown, within the Avida framework, to distinguish between co-existence of cooperators and cheaters, and exclusive persistence of cheaters in a threshold-based manner (3). We have not explicitly observed such cooperator-cheater dichotomy in our results (due perhaps to the fact that our simulation design was not optimized to ecological scales). However, we do see that these two factors regulate fitness in complex and distinct ways under high and low mutation rates. For example, incidence of runs that result in acquisition of the lowest and highest fitness genotypes (through increase in replication rate) show little variation on environmental input change, when mutation rates are low. It must be admitted though that higher values of the environmental inputs do favor greater frequencies of reproductively efficient high fitness genotypes (specifically L3 but also L2) over relatively inefficient counterparts (L1). This suggests that an appropriate change in environmental parameters is permissive to genotypes acquiring beneficial mutations even under low rates.

When the mutation rate is high, environmental variation has no effect on the lowest fitness genotypic frequency (H0) and its effect on high fitness genotypic frequency (peaks H2 and H3) is non-monotonic and ambiguous. Only the frequency of suboptimal fitness genotype (H1) acquired through cooperation under high mutation rates, shows a direct correlation with population size inputs. The interplay between mutation rates and effective population size on population dynamics on an adaptive landscape has been demonstrated using RNA folding by Vahdati and coworkers (25). Additionally, we observe that fitness values increase with greater resource levels, but only in the background of high population size. These observations establish the importance of the interdependence between the three input variables we have chosen in the study and show how fitness levels, as well as the mode of fitness acquisition are nuanced multidimensional outputs of the genotype and the environment within which it evolves.

High mutation rates also allow the evolution of higher fitness through cooperation of organisms (peak H1). Even in this case, messaging instructions that allow for cooperation appear in other genotypes but their contribution to fitness for such genotypes is minimal, compared to H1 genotype whose fitness depends on such instructions. These observations are consistent with the robust-yet-fragile hypothesis extended for complex adaptive systems, wherein fitness of genotypes could evolve robustness against a contingency while remaining sensitive to others (26). It is pertinent to wonder why cooperation evolves in the first place when it neither achieves very high fitness values nor is it robust to mutations in the messaging instructions, unlike other peak genotypes that arise under high mutation rates. One answer to this comes from our analysis that populations dominated by the cooperative genotypes tend to have more genetic heterogeneity than the other three peaks: by keeping its fitness levels low, a greater exploration of spatially coincident genetic variation can be maintained within populations without clonal interference.

Whereas our evolutionary framework captures sequence evolution and its correlation with evolution of genotypes fairly well, we wish to emphasize a few notable limitations. An important one is the absence of a developmental time scale and phenotypic (morphological and behavioral) traits associated with it. Incorporation of this time scale would allow the testing of the effects of nonadaptive plasticity of the phenotype on its evolution within populations (27,28) and in a broader context the effect of development on the evolution of phenotype (29–32). Even if the developmental time scale is not explicitly incorporated in our framework, our demonstration of a cooperative mode of evolution under high mutation rates and environment-permissive conditions provide insights into how multicellular organization could have emerged from unicellular life-histories (33,34). In fact, efforts to explain evolutionary bursts within phyla or rapid speciation predict higher rates of mutation or genomic evolution as causative to their occurrence (35–37).

The framework also implicitly assumes that the response to environmental changes is directly coded if at all within the genome, disallowing the potential to test environmental effects on epigenetic mechanisms contributing to phenotypic evolution. The evolutionary process in Avida can thus affect changes that take place at the scale of a single organismal lifetime to a greater extent than it does for real biological systems. We have also not allowed our simulations to run across huge population sizes: doing so may allow us to witness the dynamics of coexistent genotypic mutants which we seldom see in our runs. However, our motivation in this study was to study evolution within smaller asexual populations, such as in the case of transitory metastatic niches of dividing cancer cells, and derive fundamental principles for the their behaviors over long time scales (38). It has not escaped our notice that high mutation rates under appropriate environmental contexts engender the evolution of plastic, heterogeneous and cooperative populations, properties increasingly being attributed as fundamental to malignant neoplasms (39,40). A more rigorous comparison between these two systems will be undertaken in the near future.

We conclude by pointing out that our observations reinforce the fact that specific environmental variables play an important role in determining both the variety and the probability of evolutionary outcomes that arise within finite populations. In the background of a higher rate of mutation, such variables even facilitate the evolution of cooperation within genetically heterogeneous organismal ensembles.

## Acknowledgements

This work was supported by the Wellcome Trust/DBT India Alliance Fellowship/Grant [Grant Number IA/I/17/2/503312] awarded to Ramray Bhat. RB also acknowledges support from DBT [BT/PR26526/GET/119/92/2017]. BK acknowledges KVPY for the scholarship.

## Footnotes

1 Note that the measurement of merit still takes place in a test CPU and thus cannot account for ecological information sharing as is seen in peak 1 genomes.

## Notes

### Competing Interest Statement

The authors have declared no competing interest.

