## Supplementary Figures/Tables for "Distinct evolutionary trajectories in asexual populations through an interplay of their size, resource availability and mutation rates"

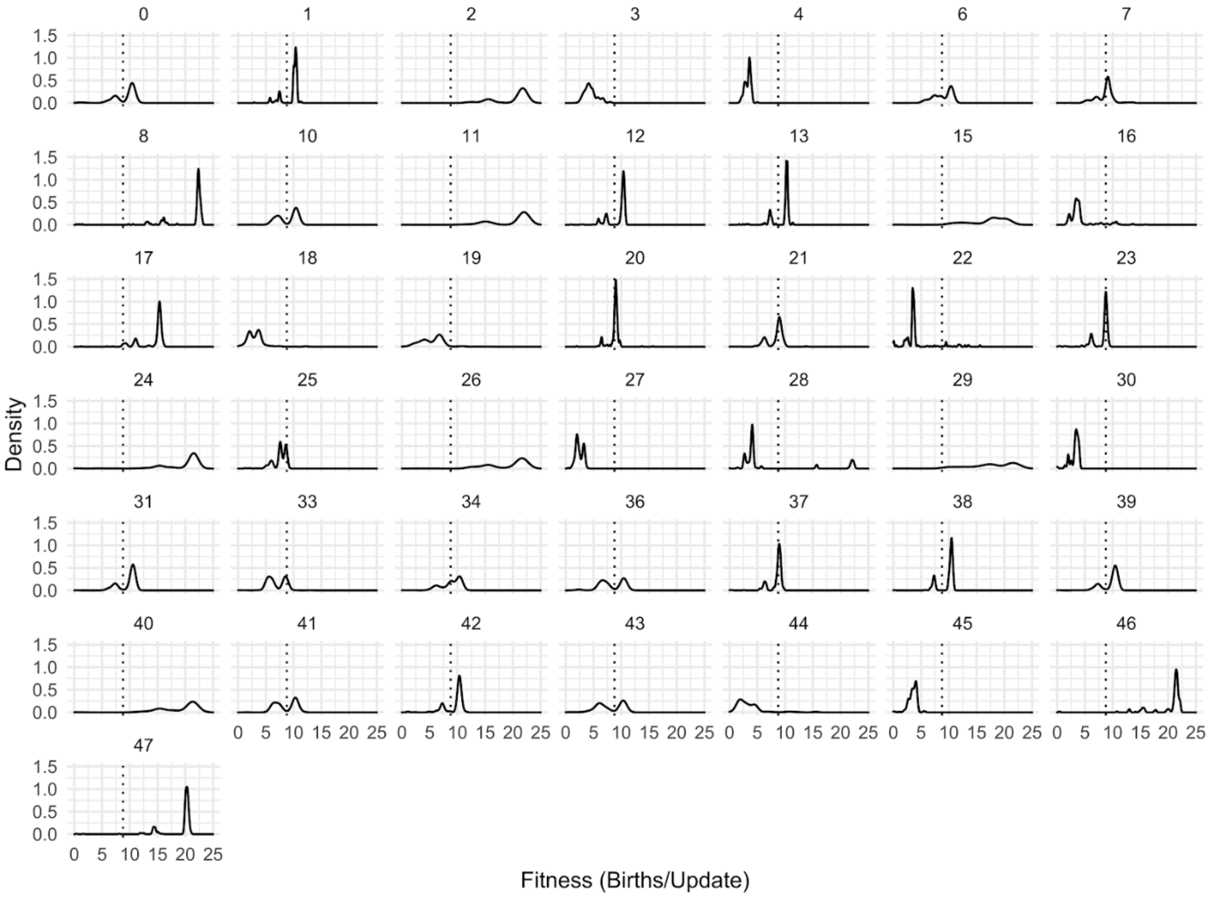

*Supplementary Figure 1. Fitness distributions for all populations obtained at a high mutation rate at a moderate resource availability (100k) and a high population size (500).*

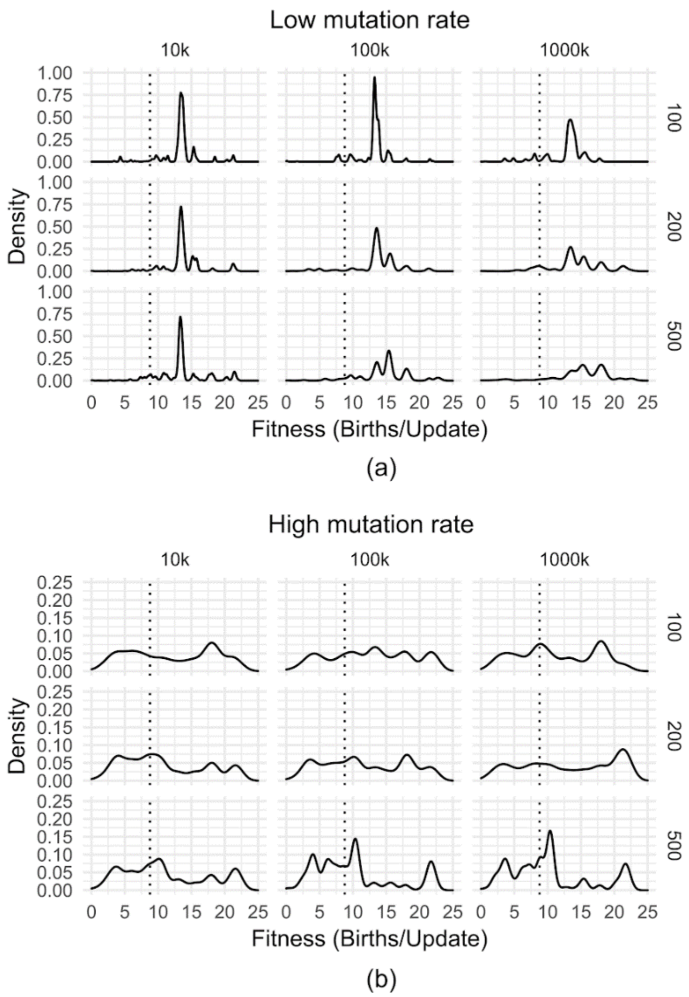

(a)

(b)

*Supplementary Figure 2. Fitness distributions of genotypes obtained for different values of resource availability and population size - (a) at a low mutation rate, and (b) at a high mutation rate.*

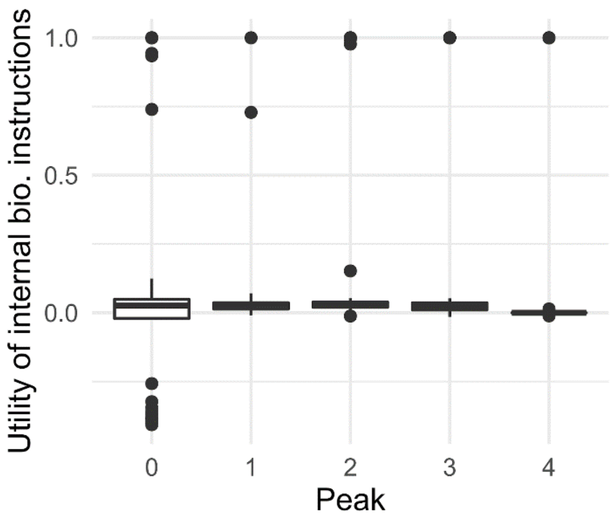

*Supplementary Figure 3. Marginal utility of internal biological instructions for genotypes belonging to different peaks at the low mutation rate (L0-L4).*

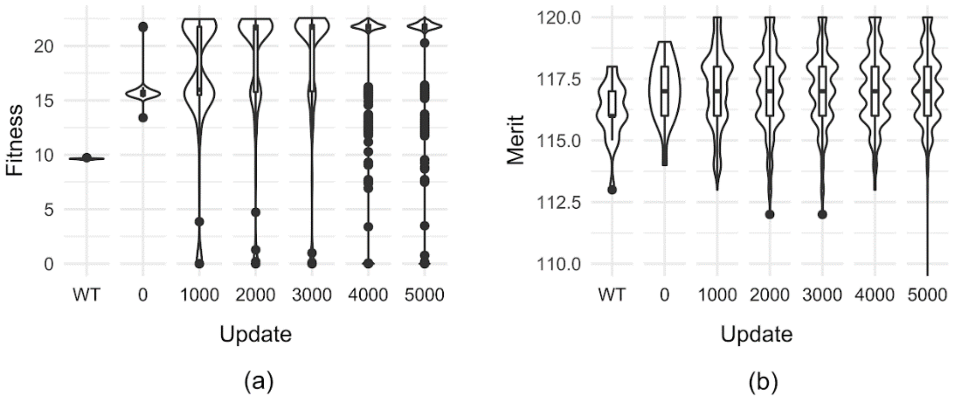

*Supplementary Figure 4. (a) Fitness and (b) merit evolution when recipient genomes sampled from peak L0 are transplanted with peak L4 copy-loop and evolved for different numbers of updates. Only viable recipient genomes are chosen (Viability ~ 19%). Updates on the x-axis is the evolutionary time for which the hybrid genotypes were evolved. "WT" denotes these measures for the recipient genomes without copy-loop replacement.*

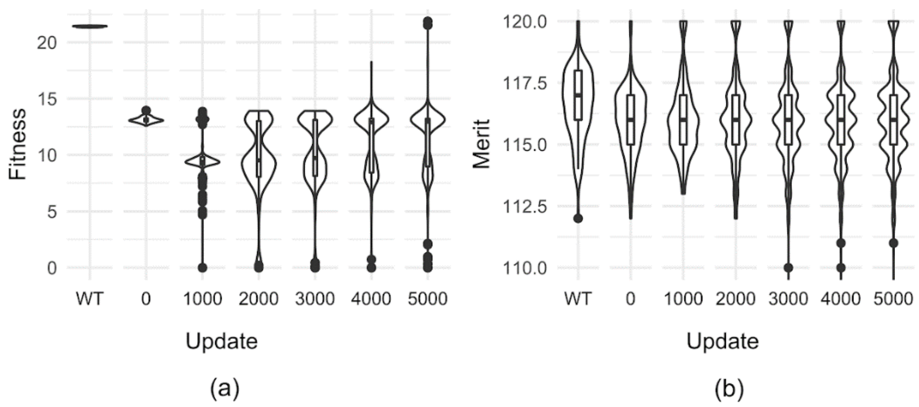

*Supplementary Figure 5. (a) Fitness and (b) merit evolution when recipient genomes sampled from peak L4 are transplanted with peak L0 copy-loop and evolved. Only viable recipient genomes are chosen (Viability ~ 100%). Updates on the x-axis is the evolutionary time for which the hybrid genotypes were evolved. "WT" denotes these measures for the recipient genomes without copy-loop replacement.*

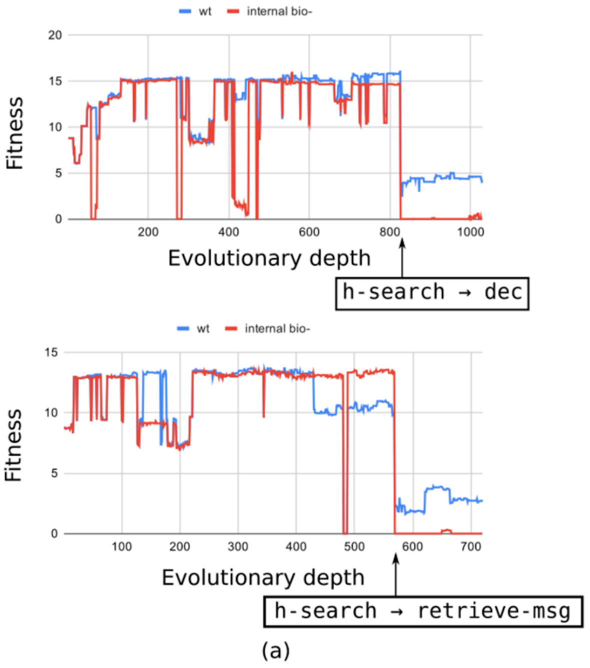

*Supplementary Figure 6. (a) Two representative curves showing the temporal fitness variation of genome lineages from peak L0. The major fitness decline is shown by arrows indicating the mutation that happens at these events. Note that the internal biological instruction knockouts immediately fall to a fitness of zero at these steps. The red line shows the fitness for the lineage after internal biological instructions are knocked out. The difference in the wild type and this fitness is seen only after the loss of h-search indicating the internalization is pre-existent (b) Internal biological instructions generate fitness to a large extent only for genomes from peak L0.*

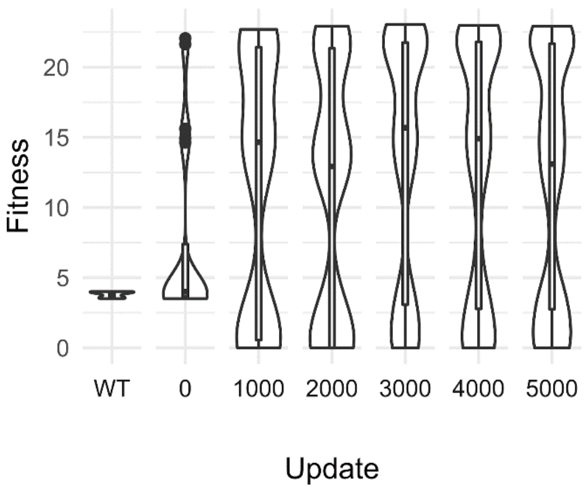

*Supplementary Figure 7. Fitness evolution when recipient genomes sampled from peak H0 are transplanted with H3 copy-loop and evolved. Only viable recipient genomes are chosen (Viability ~ 20%). Updates on the x-axis is the evolutionary time for which the hybrid genotypes were evolved. "WT" denotes these measures for the recipient genomes without copy-loop replacement.*

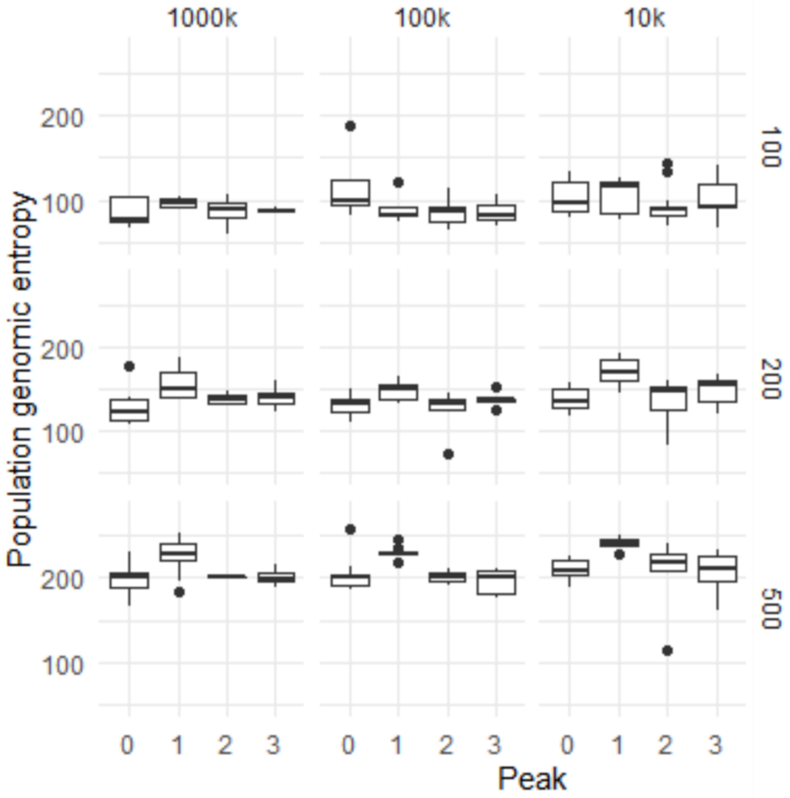

*Supplementary Figure 8. Genomic heterogeneity for populations belonging to the different peaks obtained at a high mutation rate measured by calculating the sum of per-site genomic entropies - divided by resource availability and population size.*

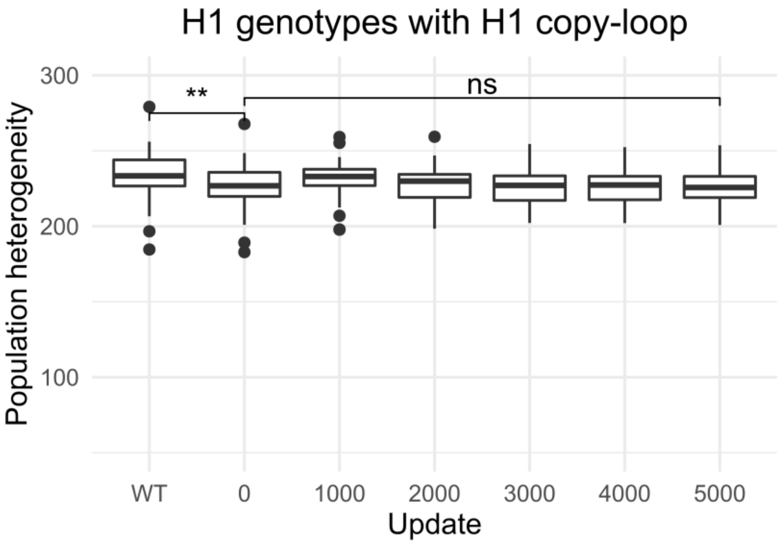

*Supplementary Figure 9. Evolution of population heterogeneity when populations dominated by H1 genotypes are transplanted with a single H1 copy-loop. (Populations evolved under maximum size 500 and 1000k resource availability)*

| | Lower | Upper |
|---|---|---|
| **L0** | 9.4508777 | 9.8534563 |
| **L1** | 13.2554407 | 13.6580193 |
| **L2** | 15.1577207 | 15.5602993 |
| **L3** | 17.8404207 | 18.2429993 |
| **L4** | 21.2059907 | 21.6085693 |
| **H0** | 3.2196089 | 4.3603911 |
| **H1** | 9.7764489 | 10.9172311 |
| **H2** | 17.4385589 | 18.5793411 |
| **H3** | 21.0846689 | 22.2254511 |

*Supplementary Table 1. Peak fitness ranges used for classification of genotypes. This range is calculated by taking a window equal in size to twice the distribution bandwidth around the maxima obtained from the cumulative fitness distribution.*

| LOW MUT. RATE | | NOT | NAND | AND | ORN | OR | ANDN | NOR | XOR | EQU |
|---|---|---|---|---|---|---|---|---|---|---|
| Peak 0 | LQ | 0 | 0 | 0 | 0 | 0 | 0 | 0 | 0 | 0 |
| | MED | 0 | 1 | 0 | 2 | 0 | 0 | 0 | 0 | 0 |
| | UQ | 0 | 6 | 11 | 20 | 12 | 3 | 2 | 0 | 0 |
| Peak 1 | LQ | 0 | 0 | 0 | 0 | 0 | 0 | 0 | 0 | 0 |
| | MED | 0 | 1 | 0 | 7 | 0 | 0 | 0 | 0 | 0 |
| | UQ | 0 | 2 | 1 | 12 | 6 | 10 | 0 | 0 | 0 |
| Peak 2 | LQ | 0 | 1 | 0 | 0 | 0 | 0 | 0 | 0 | 0 |
| | MED | 0 | 1 | 0 | 4 | 0 | 0 | 0 | 0 | 0 |
| | UQ | 0 | 3 | 4 | 14 | 11 | 9 | 0 | 0 | 0 |
| Peak 3 | LQ | 0 | 1 | 0 | 0 | 0 | 0 | 0 | 0 | 0 |
| | MED | 0 | 1 | 0 | 4 | 0 | 7 | 0 | 0 | 0 |
| | UQ | 0 | 1 | 8 | 14 | 1 | 20 | 0 | 0 | 0 |
| Peak 4 | LQ | 0 | 0 | 0 | 0 | 0 | 0 | 0 | 0 | 0 |
| | MED | 0 | 1 | 0 | 0 | 0 | 2 | 0 | 0 | 0 |
| | UQ | 1 | 2 | 7 | 9 | 11 | 11 | 2 | 0 | 0 |

| HIGH MUT. RATE | | NOT | NAND | AND | ORN | OR | ANDN | NOR | XOR | EQU |
|---|---|---|---|---|---|---|---|---|---|---|
| Peak 0 | LQ | 0 | 0 | 0 | 0 | 0 | 0 | 0 | 0 | 0 |
| | MED | 0 | 0 | 0 | 0 | 0 | 0 | 0 | 0 | 0 |
| | UQ | 0 | 7 | 0 | 9 | 60 | 0 | 60 | 0 | 0 |
| Peak 1 | LQ | 0 | 0 | 0 | 0 | 0 | 0 | 0 | 0 | 0 |
| | MED | 0 | 0 | 0 | 0 | 0 | 0 | 0 | 0 | 0 |
| | UQ | 0 | 0 | 0 | 1 | 2 | 0 | 117 | 0 | 0 |
| Peak 2 | LQ | 0 | 0 | 0 | 0 | 0 | 0 | 0 | 0 | 0 |
| | MED | 0 | 0 | 0 | 0 | 0 | 0 | 0 | 0 | 0 |
| | UQ | 0 | 0 | 0 | 1 | 1 | 0 | 12 | 0 | 0 |
| Peak 3 | LQ | 0 | 0 | 0 | 0 | 0 | 0 | 0 | 0 | 0 |
| | MED | 0 | 0 | 0 | 0 | 0 | 0 | 0 | 0 | 0 |
| | UQ | 0 | 0 | 0 | 1 | 2 | 0 | 0 | 0 | 0 |

*Supplementary Table 2. Lower quartile (LQ), upper quartile (UQ) and median values of task instances for each task for genomes from low and high mutation rates. (Data for Figure 2c and Figure 3c)*

| | | Fraction of all genotypes | |
|---|---|---|---|
| **Resources** | **Population** | **Peak 2** | **Peak 3** |
| 10,000 | 100 | 0.056064 | 0 |
| 1,000,000 | 500 | 0.130492 | 0.142857 |

*Supplementary Table 3. Proportion of peak L2 and L3 genotypes at a low mutation rate at the two extreme values of resource levels and population size.*

| L0 | | | L1 | | | L2 | | | L3 | | | L4 | | |
|---|---|---|---|---|---|---|---|---|---|---|---|---|---|---|
| **contrast** | **t.ratio** | **p.value** | **contrast** | **t.ratio** | **p.value** | **contrast** | **t.ratio** | **p.value** | **contrast** | **t.ratio** | **p.value** | **contrast** | **t.ratio** | **p.value** |
| 100 10 - 200 10 | -42.624 | <.0001 | 100 10 - 200 10 | -5.506 | <.0001 | 100 10 - 200 10 | 19.52 | <.0001 | 100 10 - 200 10 | 22.91 | <.0001 | 100 10 - 200 10 | 12.056 | <.0001 |
| 100 10 - 500 10 | -13.519 | <.0001 | 100 10 - 500 10 | -42.206 | <.0001 | 100 10 - 500 10 | 73.689 | <.0001 | 100 10 - 500 10 | 53.718 | <.0001 | 100 10 - 500 10 | -12.158 | <.0001 |
| 100 10 - 100 100 | -18.506 | <.0001 | 100 10 - 100 100 | -26.149 | <.0001 | 100 10 - 100 100 | 47.104 | <.0001 | 100 10 - 100 100 | 7.444 | <.0001 | 100 10 - 100 100 | -4.73 | <.0001 |
| 100 10 - 200 100 | 13.834 | <.0001 | 100 10 - 200 100 | -29.814 | <.0001 | 100 10 - 200 100 | 49.31 | <.0001 | 100 10 - 200 100 | 29.479 | <.0001 | 100 10 - 200 100 | 7.065 | <.0001 |
| 100 10 - 500 100 | -33.273 | <.0001 | 100 10 - 500 100 | 12.251 | <.0001 | 100 10 - 500 100 | -0.781 | 0.4346 | 100 10 - 500 100 | 39.182 | <.0001 | 100 10 - 500 100 | -0.381 | 0.7031 |
| 100 10 - 100 1000 | -29.63 | <.0001 | 100 10 - 100 1000 | -22.442 | <.0001 | 100 10 - 100 1000 | 47.697 | <.0001 | 100 10 - 100 1000 | 65.739 | <.0001 | 100 10 - 100 1000 | -3.411 | 0.0007 |
| 100 10 - 200 1000 | -26.345 | <.0001 | 100 10 - 200 1000 | -10.226 | <.0001 | 100 10 - 200 1000 | 18.565 | <.0001 | 100 10 - 200 1000 | 28.451 | <.0001 | 100 10 - 200 1000 | -18.065 | <.0001 |
| 100 10 - 500 1000 | 4.974 | <.0001 | 100 10 - 500 1000 | -14.969 | <.0001 | 100 10 - 500 1000 | 24.43 | <.0001 | 100 10 - 500 1000 | 0.392 | 0.6951 | 100 10 - 500 1000 | 21.051 | <.0001 |
| 200 10 - 500 10 | 29.105 | <.0001 | 200 10 - 500 10 | -36.7 | <.0001 | 200 10 - 500 10 | 54.169 | <.0001 | 200 10 - 500 10 | 30.808 | <.0001 | 200 10 - 500 10 | -24.213 | <.0001 |
| 200 10 - 100 100 | 24.119 | <.0001 | 200 10 - 100 100 | -20.644 | <.0001 | 200 10 - 100 100 | 27.584 | <.0001 | 200 10 - 100 100 | -15.466 | <.0001 | 200 10 - 100 100 | -16.785 | <.0001 |
| 200 10 - 200 100 | 56.458 | <.0001 | 200 10 - 200 100 | -24.308 | <.0001 | 200 10 - 200 100 | 29.79 | <.0001 | 200 10 - 200 100 | 6.57 | <.0001 | 200 10 - 200 100 | -4.99 | <.0001 |
| 200 10 - 500 100 | 9.351 | <.0001 | 200 10 - 500 100 | 17.756 | <.0001 | 200 10 - 500 100 | -20.301 | <.0001 | 200 10 - 500 100 | 16.272 | <.0001 | 200 10 - 500 100 | -12.437 | <.0001 |
| 200 10 - 100 1000 | 12.994 | <.0001 | 200 10 - 100 1000 | -16.936 | <.0001 | 200 10 - 100 1000 | 28.177 | <.0001 | 200 10 - 100 1000 | 42.829 | <.0001 | 200 10 - 100 1000 | -15.466 | <.0001 |
| 200 10 - 200 1000 | 16.279 | <.0001 | 200 10 - 200 1000 | -4.72 | <.0001 | 200 10 - 200 1000 | -0.955 | 0.3394 | 200 10 - 200 1000 | 5.541 | <.0001 | 200 10 - 200 1000 | -30.121 | <.0001 |
| 200 10 - 500 1000 | 47.598 | <.0001 | 200 10 - 500 1000 | -9.464 | <.0001 | 200 10 - 500 1000 | 4.91 | <.0001 | 200 10 - 500 1000 | -22.518 | <.0001 | 200 10 - 500 1000 | 8.996 | <.0001 |
| 500 10 - 100 100 | -4.987 | <.0001 | 500 10 - 100 100 | 16.057 | <.0001 | 500 10 - 100 100 | -26.585 | <.0001 | 500 10 - 100 100 | -46.274 | <.0001 | 500 10 - 100 100 | 7.428 | <.0001 |
| 500 10 - 200 100 | 27.353 | <.0001 | 500 10 - 200 100 | 12.392 | <.0001 | 500 10 - 200 100 | -24.379 | <.0001 | 500 10 - 200 100 | -24.239 | <.0001 | 500 10 - 200 100 | 19.223 | <.0001 |
| 500 10 - 500 100 | -19.754 | <.0001 | 500 10 - 500 100 | 54.456 | <.0001 | 500 10 - 500 100 | -74.47 | <.0001 | 500 10 - 500 100 | -14.536 | <.0001 | 500 10 - 500 100 | 11.776 | <.0001 |
| 500 10 - 100 1000 | -16.112 | <.0001 | 500 10 - 100 1000 | 19.764 | <.0001 | 500 10 - 100 1000 | -25.992 | <.0001 | 500 10 - 100 1000 | 12.021 | <.0001 | 500 10 - 100 1000 | 8.747 | <.0001 |
| 500 10 - 200 1000 | -12.826 | <.0001 | 500 10 - 200 1000 | 31.98 | <.0001 | 500 10 - 200 1000 | -55.124 | <.0001 | 500 10 - 200 1000 | -25.267 | <.0001 | 500 10 - 200 1000 | -5.907 | <.0001 |
| 500 10 - 500 1000 | 18.493 | <.0001 | 500 10 - 500 1000 | 27.236 | <.0001 | 500 10 - 500 1000 | -49.259 | <.0001 | 500 10 - 500 1000 | -53.326 | <.0001 | 500 10 - 500 1000 | 33.209 | <.0001 |
| 100 100 - 200 100 | 32.34 | <.0001 | 100 100 - 200 100 | -3.664 | 0.0002 | 100 100 - 200 100 | 2.206 | 0.0274 | 100 100 - 200 100 | 22.035 | <.0001 | 100 100 - 200 100 | 11.795 | <.0001 |
| 100 100 - 500 100 | -14.768 | <.0001 | 100 100 - 500 100 | 38.4 | <.0001 | 100 100 - 500 100 | -47.885 | <.0001 | 100 100 - 500 100 | 31.738 | <.0001 | 100 100 - 500 100 | 4.349 | <.0001 |
| 100 100 - 100 1000 | -11.125 | <.0001 | 100 100 - 100 1000 | 3.708 | 0.0002 | 100 100 - 100 1000 | 0.593 | 0.5531 | 100 100 - 100 1000 | 58.295 | <.0001 | 100 100 - 100 1000 | 1.319 | 0.1872 |
| 100 100 - 200 1000 | -7.839 | <.0001 | 100 100 - 200 1000 | 15.923 | <.0001 | 100 100 - 200 1000 | -28.539 | <.0001 | 100 100 - 200 1000 | 21.007 | <.0001 | 100 100 - 200 1000 | -13.335 | <.0001 |
| 100 100 - 500 1000 | 23.48 | <.0001 | 100 100 - 500 1000 | 11.18 | <.0001 | 100 100 - 500 1000 | -22.674 | <.0001 | 100 100 - 500 1000 | -7.052 | <.0001 | 100 100 - 500 1000 | 25.781 | <.0001 |
| 200 100 - 500 100 | -47.107 | <.0001 | 200 100 - 500 100 | 42.064 | <.0001 | 200 100 - 500 100 | -50.091 | <.0001 | 200 100 - 500 100 | 9.703 | <.0001 | 200 100 - 500 100 | -7.447 | <.0001 |
| 200 100 - 100 1000 | -43.464 | <.0001 | 200 100 - 100 1000 | 7.372 | <.0001 | 200 100 - 100 1000 | -1.613 | 0.1069 | 200 100 - 100 1000 | 36.26 | <.0001 | 200 100 - 100 1000 | -10.476 | <.0001 |
| 200 100 - 200 1000 | -40.179 | <.0001 | 200 100 - 200 1000 | 19.587 | <.0001 | 200 100 - 200 1000 | -30.745 | <.0001 | 200 100 - 200 1000 | -1.028 | 0.3039 | 200 100 - 200 1000 | -25.131 | <.0001 |
| 200 100 - 500 1000 | -8.86 | <.0001 | 200 100 - 500 1000 | 14.844 | <.0001 | 200 100 - 500 1000 | -24.879 | <.0001 | 200 100 - 500 1000 | -29.087 | <.0001 | 200 100 - 500 1000 | 13.986 | <.0001 |
| 500 100 - 100 1000 | 3.643 | 0.0003 | 500 100 - 100 1000 | -34.692 | <.0001 | 500 100 - 100 1000 | 48.478 | <.0001 | 500 100 - 100 1000 | 26.557 | <.0001 | 500 100 - 100 1000 | -3.03 | 0.0025 |
| 500 100 - 200 1000 | 6.928 | <.0001 | 500 100 - 200 1000 | -22.477 | <.0001 | 500 100 - 200 1000 | 19.346 | <.0001 | 500 100 - 200 1000 | -10.731 | <.0001 | 500 100 - 200 1000 | -17.684 | <.0001 |
| 500 100 - 500 1000 | 38.247 | <.0001 | 500 100 - 500 1000 | -27.22 | <.0001 | 500 100 - 500 1000 | 25.211 | <.0001 | 500 100 - 500 1000 | -38.79 | <.0001 | 500 100 - 500 1000 | 21.432 | <.0001 |
| 100 1000 - 200 1000 | 3.285 | 0.001 | 100 1000 - 200 1000 | 12.216 | <.0001 | 100 1000 - 200 1000 | -29.132 | <.0001 | 100 1000 - 200 1000 | -37.288 | <.0001 | 100 1000 - 200 1000 | -14.654 | <.0001 |
| 100 1000 - 500 1000 | 34.604 | <.0001 | 100 1000 - 500 1000 | 7.472 | <.0001 | 100 1000 - 500 1000 | -23.267 | <.0001 | 100 1000 - 500 1000 | -65.347 | <.0001 | 100 1000 - 500 1000 | 24.462 | <.0001 |
| 200 1000 - 500 1000 | 31.319 | <.0001 | 200 1000 - 500 1000 | -4.743 | <.0001 | 200 1000 - 500 1000 | 5.866 | <.0001 | 200 1000 - 500 1000 | -28.059 | <.0001 | 200 1000 - 500 1000 | 39.116 | <.0001 |

*Supplementary Table 4. Pairwise significance values calculated for environmental inputs from Analysis of Variance of Aligned Rank Transformed Data (ART ANOVA) for the peak fractions observed at a low mutation rate. (Factors with p>0.05 are highlighted in red, listed as pop1 res1 – pop2 res2 where pop* and res* are maximum population size and resource availability)*

| H0 | | | H1 | | | H2 | | | H3 | | |
|---|---|---|---|---|---|---|---|---|---|---|---|
| **contrast** | **t.ratio** | **p.value** | **contrast** | **t.ratio** | **p.value** | **contrast** | **t.ratio** | **p.value** | **contrast** | **t.ratio** | **p.value** |
| 100 10 - 200 10 | -5.046 | <.0001 | 100 10 - 200 10 | -7.862 | <.0001 | 100 10 - 200 10 | 23.78 | <.0001 | 100 10 - 200 10 | 14.161 | <.0001 |
| 100 10 - 500 10 | 17.046 | <.0001 | 100 10 - 500 10 | 17.308 | <.0001 | 100 10 - 500 10 | -13.387 | <.0001 | 100 10 - 500 10 | -4.373 | <.0001 |
| 100 10 - 100 100 | 9.506 | <.0001 | 100 10 - 100 100 | 4.299 | <.0001 | 100 10 - 100 100 | 34.009 | <.0001 | 100 10 - 100 100 | -22.083 | <.0001 |
| 100 10 - 200 100 | 4.365 | <.0001 | 100 10 - 200 100 | 0.001 | 0.9992 | 100 10 - 200 100 | -26.448 | <.0001 | 100 10 - 200 100 | 35.904 | <.0001 |
| 100 10 - 500 100 | -2.326 | 0.02 | 100 10 - 500 100 | 6.49 | <.0001 | 100 10 - 500 100 | 8.805 | <.0001 | 100 10 - 500 100 | -0.633 | 0.5268 |
| 100 10 - 100 1000 | 0.993 | 0.3207 | 100 10 - 100 1000 | 6.126 | <.0001 | 100 10 - 100 1000 | -18.106 | <.0001 | 100 10 - 100 1000 | 36.474 | <.0001 |
| 100 10 - 200 1000 | 13.056 | <.0001 | 100 10 - 200 1000 | 18.96 | <.0001 | 100 10 - 200 1000 | 22.997 | <.0001 | 100 10 - 200 1000 | -29.005 | <.0001 |
| 100 10 - 500 1000 | -2.411 | 0.0159 | 100 10 - 500 1000 | -11.724 | <.0001 | 100 10 - 500 1000 | 15.524 | <.0001 | 100 10 - 500 1000 | 14.679 | <.0001 |
| 200 10 - 500 10 | 22.092 | <.0001 | 200 10 - 500 10 | 25.17 | <.0001 | 200 10 - 500 10 | -37.167 | <.0001 | 200 10 - 500 10 | -18.534 | <.0001 |
| 200 10 - 100 100 | 14.552 | <.0001 | 200 10 - 100 100 | 12.161 | <.0001 | 200 10 - 100 100 | 10.229 | <.0001 | 200 10 - 100 100 | -36.244 | <.0001 |
| 200 10 - 200 100 | 9.411 | <.0001 | 200 10 - 200 100 | 7.863 | <.0001 | 200 10 - 200 100 | -50.227 | <.0001 | 200 10 - 200 100 | 21.743 | <.0001 |
| 200 10 - 500 100 | 2.72 | 0.0065 | 200 10 - 500 100 | 14.352 | <.0001 | 200 10 - 500 100 | -14.975 | <.0001 | 200 10 - 500 100 | -14.794 | <.0001 |
| 200 10 - 100 1000 | 6.039 | <.0001 | 200 10 - 100 1000 | 13.988 | <.0001 | 200 10 - 100 1000 | -41.886 | <.0001 | 200 10 - 100 1000 | 22.313 | <.0001 |
| 200 10 - 200 1000 | 18.102 | <.0001 | 200 10 - 200 1000 | 26.822 | <.0001 | 200 10 - 200 1000 | -0.783 | 0.4338 | 200 10 - 200 1000 | -43.166 | <.0001 |
| 200 10 - 500 1000 | 2.635 | 0.0084 | 200 10 - 500 1000 | -3.862 | 0.0001 | 200 10 - 500 1000 | -8.255 | <.0001 | 200 10 - 500 1000 | 0.518 | 0.6044 |
| 500 10 - 100 100 | -7.54 | <.0001 | 500 10 - 100 100 | -13.009 | <.0001 | 500 10 - 100 100 | 47.396 | <.0001 | 500 10 - 100 100 | -17.711 | <.0001 |
| 500 10 - 200 100 | -12.681 | <.0001 | 500 10 - 200 100 | -17.307 | <.0001 | 500 10 - 200 100 | -13.061 | <.0001 | 500 10 - 200 100 | 40.277 | <.0001 |
| 500 10 - 500 100 | -19.372 | <.0001 | 500 10 - 500 100 | -10.818 | <.0001 | 500 10 - 500 100 | 22.192 | <.0001 | 500 10 - 500 100 | 3.74 | 0.0002 |
| 500 10 - 100 1000 | -16.053 | <.0001 | 500 10 - 100 1000 | -11.182 | <.0001 | 500 10 - 100 1000 | -4.72 | <.0001 | 500 10 - 100 1000 | 40.847 | <.0001 |
| 500 10 - 200 1000 | -3.99 | 0.0001 | 500 10 - 200 1000 | 1.652 | 0.0986 | 500 10 - 200 1000 | 36.384 | <.0001 | 500 10 - 200 1000 | -24.633 | <.0001 |
| 500 10 - 500 1000 | -19.457 | <.0001 | 500 10 - 500 1000 | -29.032 | <.0001 | 500 10 - 500 1000 | 28.911 | <.0001 | 500 10 - 500 1000 | 19.052 | <.0001 |
| 100 100 - 200 100 | -5.141 | <.0001 | 100 100 - 200 100 | -4.297 | <.0001 | 100 100 - 200 100 | -60.457 | <.0001 | 100 100 - 200 100 | 57.987 | <.0001 |
| 100 100 - 500 100 | -11.832 | <.0001 | 100 100 - 500 100 | 2.191 | 0.0284 | 100 100 - 500 100 | -25.204 | <.0001 | 100 100 - 500 100 | 21.45 | <.0001 |
| 100 100 - 100 1000 | -8.513 | <.0001 | 100 100 - 100 1000 | 1.828 | 0.0676 | 100 100 - 100 1000 | -52.116 | <.0001 | 100 100 - 100 1000 | 58.557 | <.0001 |
| 100 100 - 200 1000 | 3.549 | 0.0004 | 100 100 - 200 1000 | 14.661 | <.0001 | 100 100 - 200 1000 | -11.012 | <.0001 | 100 100 - 200 1000 | -6.922 | <.0001 |
| 100 100 - 500 1000 | -11.917 | <.0001 | 100 100 - 500 1000 | -16.022 | <.0001 | 100 100 - 500 1000 | -18.485 | <.0001 | 100 100 - 500 1000 | 36.762 | <.0001 |
| 200 100 - 500 100 | -6.691 | <.0001 | 200 100 - 500 100 | 6.489 | <.0001 | 200 100 - 500 100 | 35.253 | <.0001 | 200 100 - 500 100 | -36.537 | <.0001 |
| 200 100 - 100 1000 | -3.372 | 0.0007 | 200 100 - 100 1000 | 6.125 | <.0001 | 200 100 - 100 1000 | 8.341 | <.0001 | 200 100 - 100 1000 | 0.57 | 0.5688 |
| 200 100 - 200 1000 | 8.691 | <.0001 | 200 100 - 200 1000 | 18.959 | <.0001 | 200 100 - 200 1000 | 49.445 | <.0001 | 200 100 - 200 1000 | -64.91 | <.0001 |
| 200 100 - 500 1000 | -6.776 | <.0001 | 200 100 - 500 1000 | -11.725 | <.0001 | 200 100 - 500 1000 | 41.972 | <.0001 | 200 100 - 500 1000 | -21.225 | <.0001 |
| 500 100 - 100 1000 | 3.319 | 0.0009 | 500 100 - 100 1000 | -0.364 | 0.716 | 500 100 - 100 1000 | -26.912 | <.0001 | 500 100 - 100 1000 | 37.107 | <.0001 |
| 500 100 - 200 1000 | 15.382 | <.0001 | 500 100 - 200 1000 | 12.47 | <.0001 | 500 100 - 200 1000 | 14.192 | <.0001 | 500 100 - 200 1000 | -28.372 | <.0001 |
| 500 100 - 500 1000 | -0.085 | 0.9326 | 500 100 - 500 1000 | -18.214 | <.0001 | 500 100 - 500 1000 | 6.719 | <.0001 | 500 100 - 500 1000 | 15.312 | <.0001 |
| 100 1000 - 200 1000 | 12.063 | <.0001 | 100 1000 - 200 1000 | 12.834 | <.0001 | 100 1000 - 200 1000 | 41.104 | <.0001 | 100 1000 - 200 1000 | -65.479 | <.0001 |
| 100 1000 - 500 1000 | -3.404 | 0.0007 | 100 1000 - 500 1000 | -17.85 | <.0001 | 100 1000 - 500 1000 | 33.631 | <.0001 | 100 1000 - 500 1000 | -21.795 | <.0001 |
| 200 1000 - 500 1000 | -15.466 | <.0001 | 200 1000 - 500 1000 | -30.684 | <.0001 | 200 1000 - 500 1000 | -7.473 | <.0001 | 200 1000 - 500 1000 | 43.685 | <.0001 |

*Supplementary Table 5. Pairwise significance values calculated for environmental inputs from Analysis of Variance of Aligned Rank Transformed Data (ART ANOVA) for the peak fractions observed at a high mutation rate. (Factors with p>0.05 are highlighted in red, listed as pop1 res1 – pop2 res2 where pop* and res* are maximum population size and resource availability)*
